# Urban waterfowl harbour geographically connected viral communities across Aotearoa New Zealand

**DOI:** 10.64898/2026.10.04.756485

**Authors:** Lia R. Heremia, Rebecca K. French, Rebecca M. Grimwood, Janelle R. Wierenga, Jérémy Dubrulle, Jordan T. Taylor, Jessica A. Darnley, Nicola S. Holdsworth, Jemma L. Geoghegan, Stephanie J. Waller

**Affiliations:** Department of Microbiology and Immunology, University of Otago, Dunedin 9016, Aotearoa New Zealand; Department of Zoology, University of Otago, Dunedin 9016, Aotearoa New Zealand; The Otago Peninsula Eco Restoration Alliance, Portobello, Dunedin, Aotearoa New Zealand

**Keywords:** Waterfowl, ducks, avian influenza A virus, highly pathogenic avian influenza (HPAI) virus, virome, distance decay, metagenomics

## Abstract

Urban waterfowl are important natural reservoirs of viruses that can affect wildlife, livestock and human health, yet the diversity and spatial distribution of these viral communities remain poorly understood. Here, we characterised the faecal viromes of urban waterfowl sampled from 13 locations across the length of Aotearoa (New Zealand). We identified 96 avian viruses (50 species) spanning nine viral families, including 33 previously undescribed viruses. Picornaviruses were the most widespread viral group, while several viruses were closely related to those previously identified in Australian waterfowl, indicating that some viral lineages are likely distributed across the broader Australasian region. Despite sampling locations spanning more than 1,500 km, virome composition exhibited little evidence of geographic structure, with little difference in viral diversity or community composition between sampling locations. Network analyses similarly demonstrated extensive sharing of virus species among sampling locations, indicating little geographic structuring of virome composition among urban waterfowl. In addition, we recovered the complete genome of a low-pathogenic A(H3N8) influenza A virus from mallards (*Anas platyrhynchos*) in the most southern region of Aotearoa. These findings reveal extensive viral diversity and widespread connectivity among urban waterfowl, providing new insights into viral ecology and the potential for widespread virus distribution across Aotearoa.

## Introduction

Waterfowl occupy a unique position in the ecology and evolution of viruses. Their high population densities, frequent interspecific mixing and, in many species, seasonal and long-distance movements create favourable conditions for the maintenance, evolution and dissemination of diverse viral communities (1,2). Consequently, ducks, geese and swans are recognised as important natural reservoirs for numerous viruses (2–5), including influenza A viruses (6), coronaviruses, astroviruses and paramyxoviruses (7). Many of these viruses circulate with little or no clinical disease in their avian hosts while retaining the capacity to infect other wildlife, domestic animals and, occasionally, humans (7,8). Waterfowl therefore represent an important wildlife interface for the emergence and dissemination of infectious diseases (8,9).

Anthropogenic environmental change has altered interactions between wildlife and people, increasing opportunities for pathogen transmission across species boundaries (2,10). Urban parks, artificial lakes and botanical gardens support large resident populations of waterfowl that frequently interact with humans and companion animals (7,9), creating environments conducive to viral transmission (2,7,11–13). Recent global outbreaks of highly pathogenic avian influenza A(H5N1) virus, for example, have highlighted the severe ecological and economic consequences that viruses circulating in wild birds can have following emergence into naïve wildlife populations and poultry (14–16). Despite the recognised importance of waterfowl in virus ecology (17–19) and the extensive diversity of avian influenza viruses they harbour (20), the diversity, distribution and geographic structure of their broader viral communities remain poorly understood in Aotearoa New Zealand (hereafter referred to as Aotearoa) (21,22).

Aotearoa’s geographic isolation has shaped a unique avifauna with historically limited natural immigration, while introduced ducks, geese and swans now dominate many urban and modified aquatic ecosystems (23–27). Avian influenza viruses are known to circulate extensively within these populations (20). Surveillance of mallards (*Anas platyrhynchos*) has identified hundreds of avian influenza viruses spanning 41 haemagglutinin (HA)/neuraminidase (NA) subtype combinations, with A(H4N6), A(H3N8), A(H3N2) and A(H10N3) among the most frequently detected, alongside low-pathogenicity H5 and H7 viruses (20). However, considerably less is known about the broader viromes of Aotearoa’s waterfowl. Whether geographically separated populations harbour distinct viral communities, the extent to which these communities vary across the country and whether they contain previously undescribed viruses remain largely unknown.

Here, we conducted a national scale metatranscriptomic survey of viruses in urban waterfowl across Aotearoa. We characterised the faecal viromes of ducks, geese and swans sampled from urban parks throughout the country to investigate viral diversity and geographic structure. We show that urban waterfowl harbour a remarkable diversity of viruses, including numerous previously undescribed species, while viral community composition exhibits no spatial structuring across large geographic distances. These findings reveal extensive viral diversity and no geographic differentiation among urban waterfowl populations, providing new insights into the processes shaping waterfowl viromes.

## Methods

### Faecal sample collection

Fresh faecal samples were collected from urban waterfowl inhabiting public parks and botanic gardens throughout Aotearoa. Sampling locations included Invercargill, Dunedin, Queenstown, Ōamaru, Christchurch, Nelson, Wellington, Paraparaumu, New Plymouth, Taupō, Rotorua, Tauranga and Auckland (Figure 1). Sites were selected based on the abundance of waterfowl and frequent human use. Where possible, each location was sampled on multiple occasions. Fresh faeces were collected immediately after defecation using sterile swabs, ensuring birds were not disturbed during sampling. Swabs were placed into 800 μL of DNA/RNA Shield (Zymo Research) and stored at 4°C before being transported to the University of Otago (Dunedin, Aotearoa), where samples were stored at −80°C until RNA extraction.

**Figure 1.**
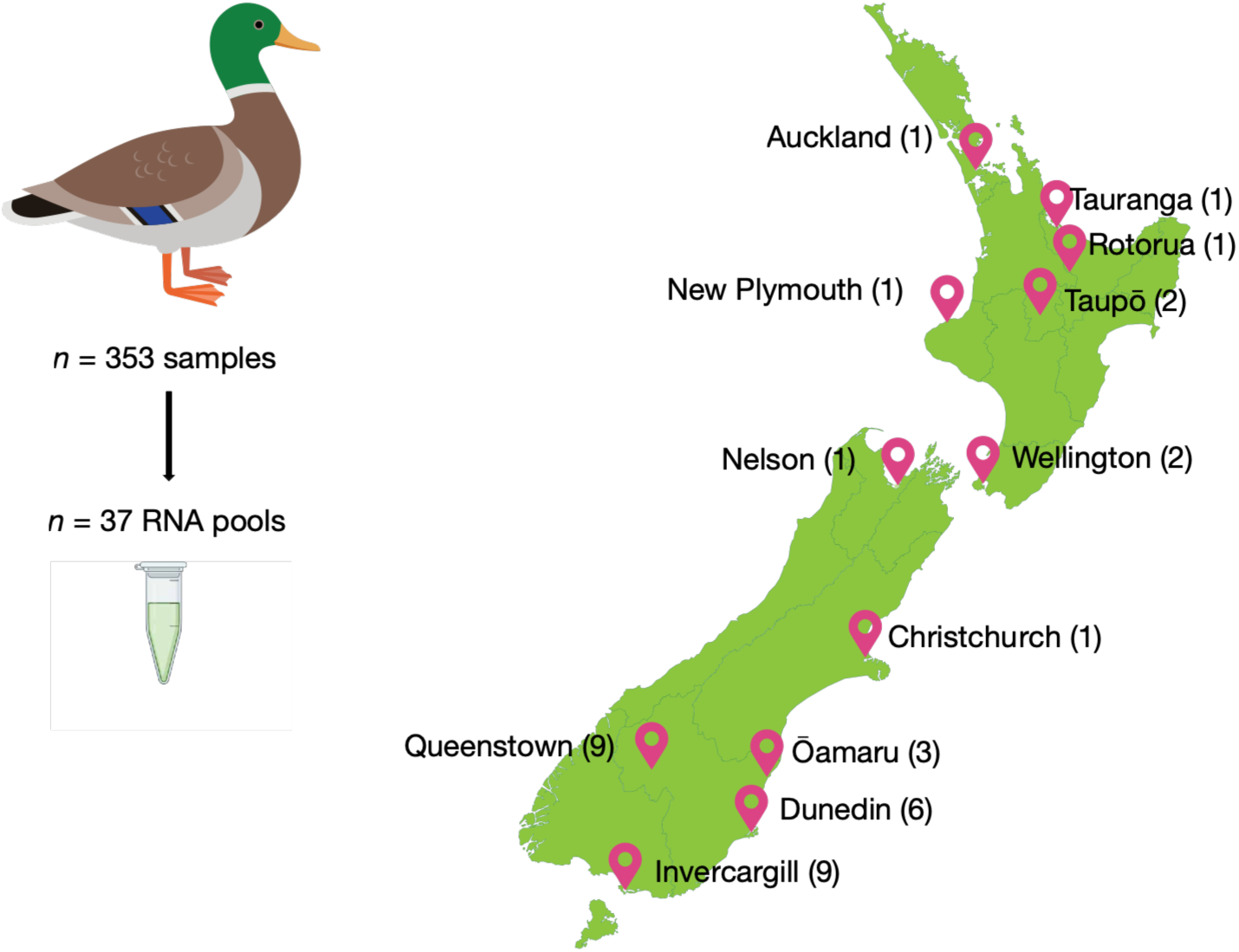
Sampling locations across Aotearoa. Numbers indicate the total RNA pools generated from each location. See Supplementary Table 1 for the number of samples per pool.

### RNA extraction

Frozen samples were thawed and transferred to ZR BashingBead Lysis tubes (0.1–0.5 mm; Zymo Research) containing 750 μL DNA/RNA Shield. Samples were homogenised using a Mini-Beadbeater-24 (BioSpec Products Inc.) for three 1.5-minute cycles with 30–45 seconds cooling intervals on ice between cycles. Total RNA was extracted using the ZymoBIOMICS MagBead RNA Kit (Zymo Research) following the manufacturer’s instructions, with additional molecular-grade ethanol washes to remove residual guanidine salts. RNA quantity and purity were assessed using a NanoDropTM One/One Microvolume UV-Vis Spectrophotometer (Thermo Fisher Scientific).

### RNA pooling and sequencing

RNA extracts were pooled in equal volumes according to sampling location and collection date, with approximately ten individual samples contributing to each pool, generating 37 pooled RNA libraries (Supplementary Table 1 and Figure 1). Sequencing libraries were prepared using the Illumina Stranded Total RNA Prep with Ribo-Zero Plus kit (Illumina) with 16 PCR cycles. Paired-end (2 × 150 bp) sequencing was performed on the NovaSeq X platform (Illumina).

### Virus identification and abundance estimation

Raw sequencing reads were quality filtered using Trimmomatic (28) with a sliding-window threshold of 5, a minimum leading and trailing base quality of 3, and a minimum read length of 25 nucleotides, and assessed with FastQC (29). Filtered reads were assembled de novo using MEGAHIT v1.2.9 (30). Assembled contigs were compared against a local copy of the NCBI non-redundant protein database using DIAMOND BLASTx (v 2.1.9) (31). Contigs were initially classified into higher taxonomic groups using the *sskingdoms* option, and non-viral matches, including bacteriophage and host-derived endogenous viral elements, were removed following manual inspection.

Putative viral contigs were further analysed in Geneious Prime (v 2026.1.1) (32) to identify open reading frames and translate contigs to protein sequences. Viral abundance was estimated by mapping quality-filtered reads back to assembled viral contigs using Bowtie2 (v.2.4.4) (33). To minimise the effects of index hopping (34), viral transcripts with abundances below 0.1% of the maximum abundance observed for that virus across all libraries were excluded from downstream analyses.

### Phylogenetic analysis

To determine the evolutionary relationships of avian viruses, maximum-likelihood phylogenetic trees were inferred for each viral family using representative reference sequences retrieved from GenBank. Conserved protein-coding regions, including the RNA-dependent RNA polymerase (RdRp) for RNA viruses and DNA polymerase for DNA viruses, were aligned using the L-INS-i algorithm in MAFFT v7.450 (35). Poorly aligned regions were removed with trimAl v1.5 (36). Phylogenetic trees were estimated in IQ-TREE (37) using the best-fitting amino acid substitution model selected by ModelFinder (38), with node support assessed using 1,000 ultrafast bootstrap replicates. Trees were visualised using FigTree (v1.4.4) (39) and Adobe Illustrator.

### Virome diversity and connectivity

All statistical analyses were performed in R (v4.6.0). Viral abundance was standardised as reads per million (RPM). Virome beta diversity was quantified using Bray-Curtis dissimilarity calculated from viral abundance matrices with the vegan package (40). Principal coordinates analysis (PCoA) was performed using the cmdscale function to visualise differences in virome composition. Libraries with no detectable avian viruses were excluded from abundance-based multivariate analyses. Permutational multivariate analysis of variance (PERMANOVA; adonis2) was used to test for differences in virome composition between the North and South Islands. Alpha diversity was quantified using the Shannon diversity index and compared between islands using Wilcoxon rank-sum tests.

To investigate spatial structure among virus species and their hosts, pairwise Bray-Curtis dissimilarity was compared with pairwise geographic distance (km) calculated between sampling locations using Google Earth (41). Viral species connectivity among sampling locations was explored using bipartite network analysis. Where viral species were identified via phylogenetic analysis, RPM values were summed across pooled libraries for each sampling location to generate a location-by-virus matrix, which was visualised using the ggbipart package (v0.1.2) (42,43). Connectance, nestedness and network specialisation were calculated using the bipartite package and network modularity was estimated using the computeModules function (44).

### Virus nomenclature

Viruses were provisionally classified as putative novel species if the most conserved protein (typically the RNA-dependent RNA polymerase or DNA polymerase) shared less than 90% amino acid identity with its closest recognised relative. Provisional common names were assigned to putative novel viruses.

## Results

### Urban waterfowl harbour diverse viromes

To characterise the diversity and connectivity of viruses circulating in urban waterfowl, we analysed 353 faecal samples collected from ducks, geese and swans across 13 urban parks and botanical gardens throughout Aotearoa between October and December 2023 (Supplementary Table 1). Samples were pooled by sampling location and collection date, generating 37 metatranscriptomic libraries for RNA sequencing. Sequencing generated an average of 153 million paired-end reads per library (range 35–271 million) (Figure 2a). De novo assembly produced between 48,963 and 498,586 contigs per library (mean = 222,605 contigs). Despite this variation, sequencing depth did not correlate with viral abundance (Figure 2b). Viral contigs represented approximately 1.49% of all assembled contigs across the 37 libraries.

**Figure 2.**
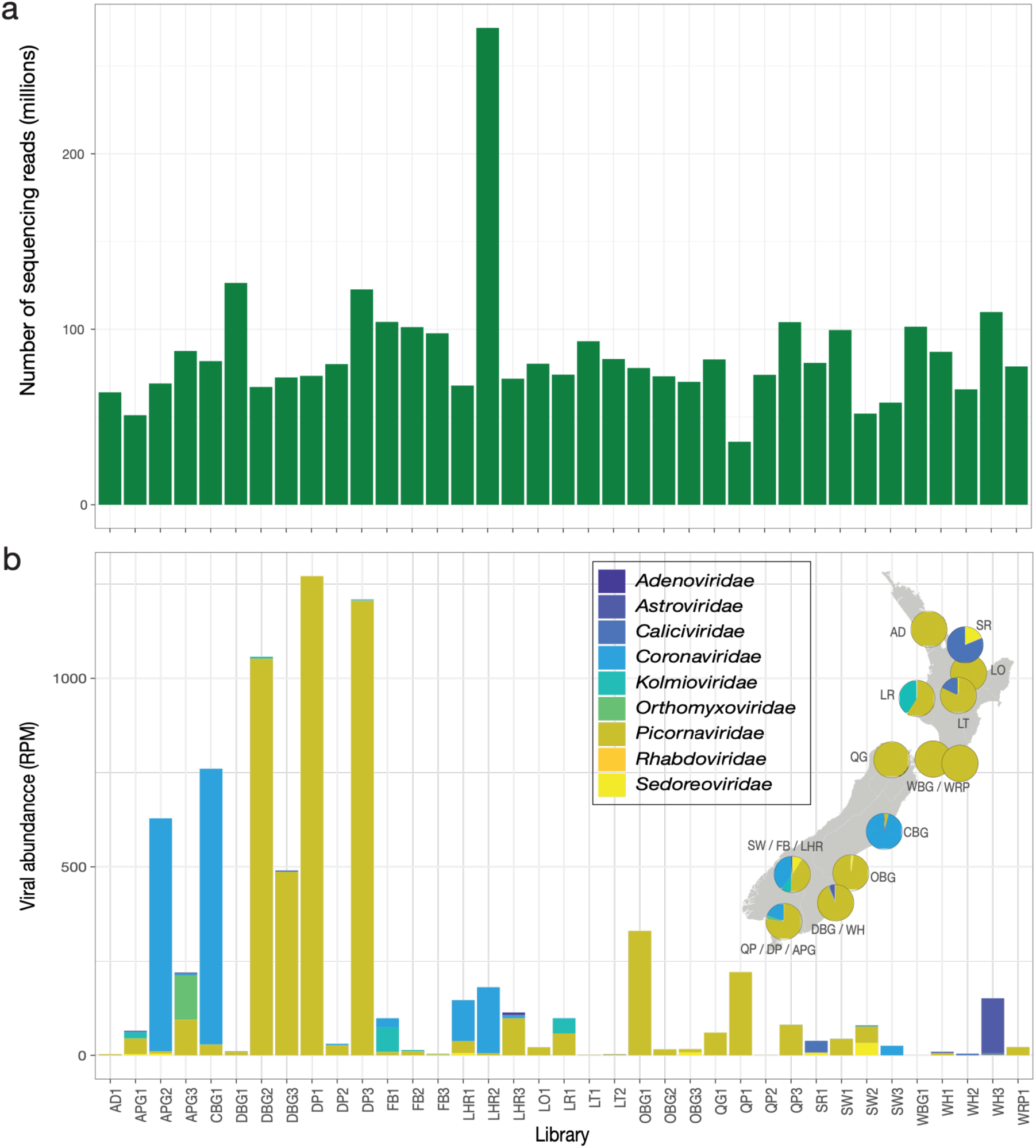
RNA sequencing depth and viral abundance across urban waterfowl libraries collected throughout Aotearoa. (**a**) Number of paired-end sequencing reads generated for each library. (**b**) Viral abundance (reads per million; RPM) across libraries, coloured by viral family. Pie charts indicate the relative abundance of viral families detected at each sampling location. The key to sampling locations is as follows: AD, Auckland; APG, DP and QP, Invercargill; DBG and WH, Dunedin; FB, LHR and SW, Queenstown; CBG, Christchurch; WRP, Paraparaumu; QG, Nelson; LR, New Plymouth; OBG, Oamaru; LT, Taupō; LO, Rotorua; WBG, Wellington; and SR, Tauranga.

Following taxonomic classification and manual curation, we identified 96 avian-associated viruses across 50 virus species, which belonged to nine viral families: *Adenoviridae*, *Astroviridae*, *Caliciviridae*, *Coronaviridae*, *Kolmioviridae*, *Orthomyxoviridae*, *Picornaviridae*, *Rhabdoviridae* and *Sedoreoviridae* (Figure 2b). Picornaviruses were the most widespread viruses, detected in 35 of the 37 libraries, followed by coronaviruses, which were detected in 15 libraries. Phylogenetic analyses based on conserved viral polymerase genes placed these viruses within established avian viral lineages while revealing substantial previously undescribed diversity.

### Urban waterfowl harbour extensive novel viral diversity

Across the 96 viruses identified, 33 represented putatively novel avian virus species, substantially expanding the known diversity of viruses associated with waterfowl in Aotearoa (Figure 3). Positive-sense RNA viruses dominated the virome. Viruses in the *Picornaviridae* exhibited the greatest diversity, comprising 37 viral transcripts distributed across multiple genera including Megrivirus, Passerivirus, Ludopivirus, Sapelovirus, Hepatovirus, Aalivirus and Harkavirus (Supplementary Table 2; Figure 3). Although many picornaviruses shared high amino acid similarity with previously described avian viruses, four highly divergent viruses clustered near the passeriviruses yet shared only 36-50% amino acid identity with recognised members of this genus, suggesting they likely represent a previously undescribed lineage.

**Figure 3.**
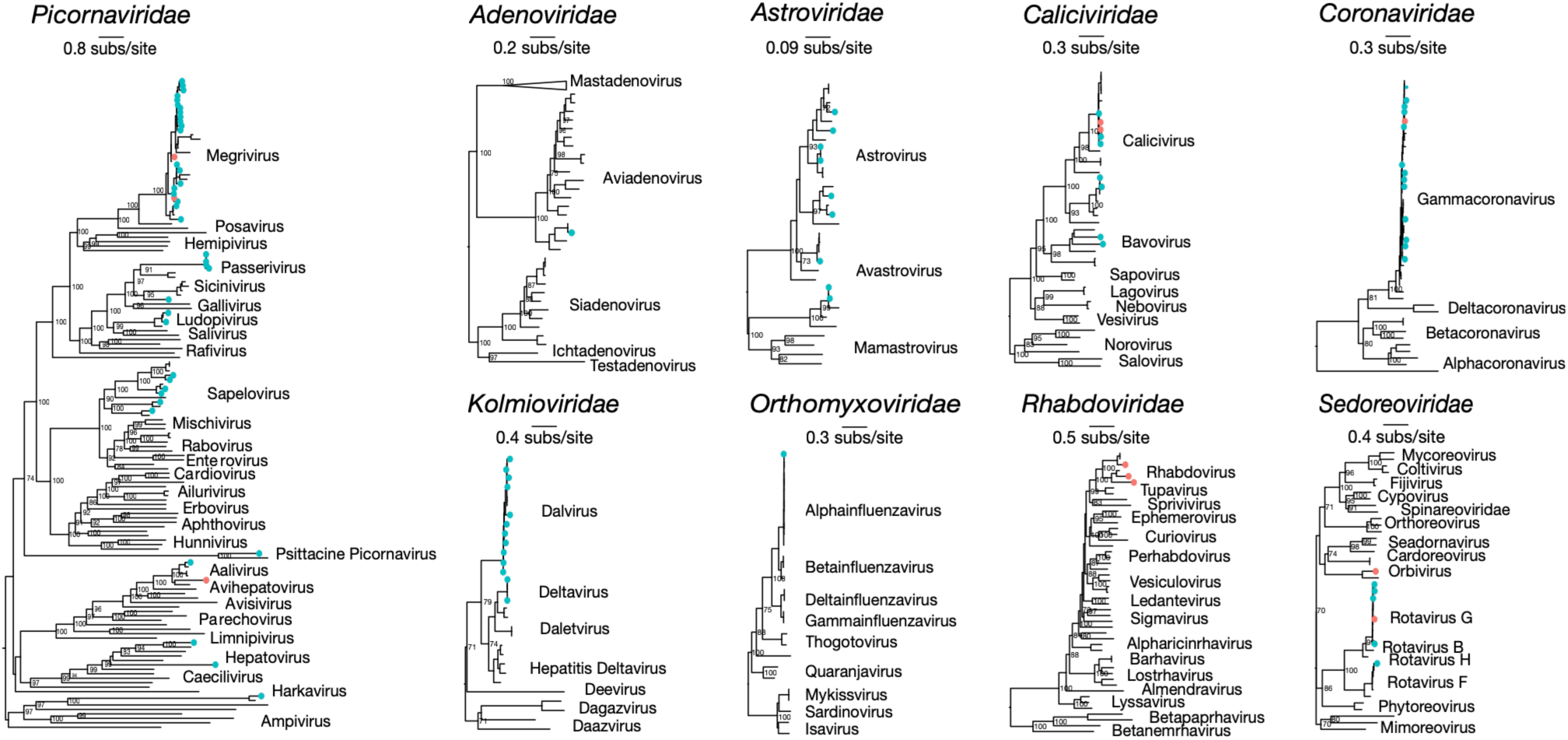
Maximum-likelihood phylogenetic trees of viruses identified in urban waterfowl. Trees were inferred from conserved viral polymerase genes for each viral family. Viruses identified in this study are indicated by coloured circles (North Island, pink; South Island, blue). Reference sequences represent recognised diversity within each viral family. Numbers at internal nodes indicate bootstrap support values. Trees are rooted at the mid-point for clarity only and branches are scaled to the number of amino acid substitutions as indicated by each scale bar.

Four novel astroviruses belonging to the genus *Avastrovirus* were identified from the lower South Island. These viruses were most closely related to astroviruses previously described from chickens and ducks in China, Poland and Australia, although each shared less than 90% amino acid identity with its nearest relative. Nine caliciviruses were also recovered, including representatives of the genera Nacovirus and Bavovirus. While most were closely related to duck caliciviruses previously reported from North America, two highly divergent bavoviruses shared only 47-51% amino acid identity with known caliciviruses from shelducks (*Tadorna tadornoides*) and mute swans (*Cygnus olor*), indicating additional undescribed diversity within this genus.

Coronaviruses were detected at four sampling locations, predominantly within the South Island (Figure 3). These viruses belonged to the genus Gammacoronavirus and exhibited greater than 99% amino acid identity to viruses recently described from Australian grey teals (*Anas gracilis*) and Australian shelducks (*Tadorna tadornoides*) (22), suggesting that closely related viral lineages circulate among waterfowl across the Australasian region.

Negative-sense RNA viruses comprised members of the *Kolmioviridae*, *Orthomyxoviridae* and *Rhabdoviridae*. Deltavirus-like sequences within the genus Dalvirus were closely related to viruses previously identified in Australian dabbling ducks (*Chenonetta jubata*) and Japanese passerines (*Pardaliparus venustulus*) (45). Three highly divergent rhabdoviruses detected in North Island waterfowl clustered with unclassified avian rhabdoviruses previously described from Australian waterfowl (21) but shared only 51-74% amino acid identity with their closest relatives, consistent with novel virus species.

Double-stranded RNA viruses were represented by the *Sedoreoviridae*, including members of the genera Rotavirus (Rotavirus F, Rotavirus G (46)) and Orbivirus (Avian-associated orbivirus 1 (47)). Most rotaviruses were closely related to viruses previously identified in Australian ducks. In addition, a double-stranded DNA virus belonging to the *Adenoviridae* was identified in samples from Queenstown. This virus clustered within the genus Aviadenovirus and shared more than 99% amino acid identity with an aviadenovirus previously detected in Pacific black ducks (*Anas superciliosa*) from Australia (47).

No highly pathogenic avian influenza viruses were detected, and sampling was conducted prior to the 2026 A(H5N1) incursion, however we recovered the complete genome of a low pathogenic H3N8 influenza A virus sampled from Anderson Park Gardens, Invercargill. This virus was designated A/mallard/NewZealand/APG3/2024(H3N8). The PB1 segment shared 100% amino acid identity with the corresponding segment of A/mallard/NewZealand/1365355/2005(H7N7) (20) (Figure 3).

### Urban waterfowl maintain a highly connected national virome

To determine whether viral communities were geographically structured, we compared virome composition at the virus species level among sampling locations spanning the length of Aotearoa (Figure 4). Pairwise virome dissimilarity was not associated with geographic distance (Mantel correlation coefficient, *r* = 0.118; *p* = 0.075) (Figure 4a). Shannon diversity did not differ significantly between the North and South Islands (Wilcoxon rank-sum test, *p* = 0.712; Figure 4b), indicating comparable levels of viral diversity between islands.

**Figure 4.**
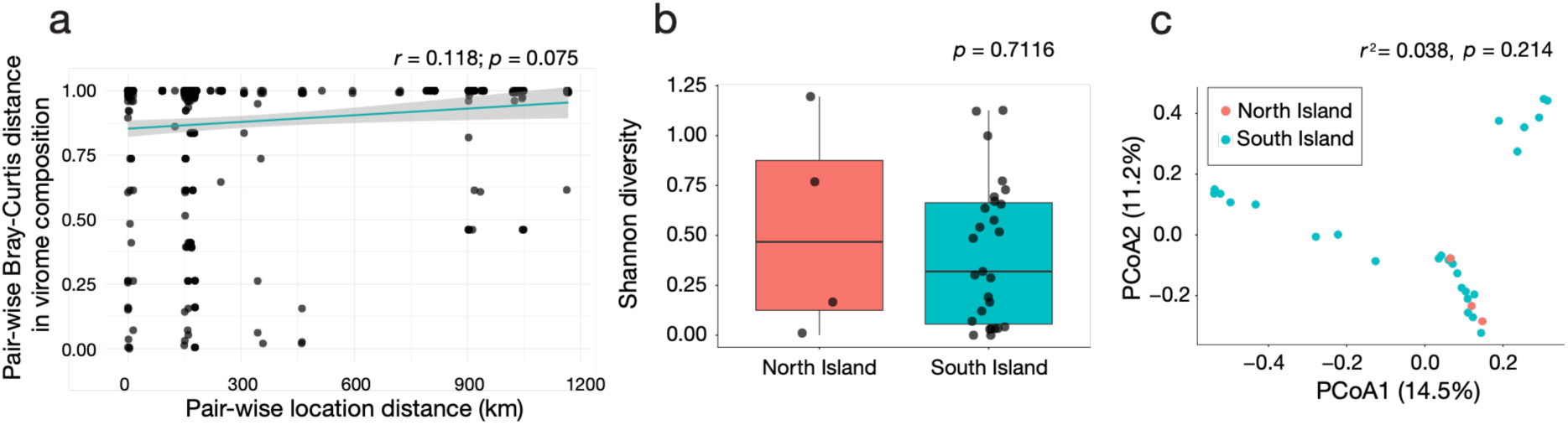
Geographic structure of urban waterfowl viromes. (**a**) Relationship between pairwise geographic distance and Bray–Curtis virome dissimilarity. (**b**) Shannon diversity of North and South Island viromes. (**c**) Principal coordinates analysis of virome composition.

Principal coordinates analysis revealed substantial overlap in virome composition among sampling locations from both islands, with no evidence of island-specific clustering (PERMANOVA, r^2^ *=* 0.038, *p* = 0.21; Figure 4c). Likewise, bipartite network analysis demonstrated that viral species were widely shared among sampling locations throughout Aotearoa rather than being restricted to individual regions (Figure 5). The network visualisation incorporated viral species abundance with connection thickness proportional to the summed RPM of each viral species at the corresponding sampling location. Note that only virus species identified through phylogenetic analysis are represented in the network. Network metrics indicated relatively low geographic compartmentalisation of the virome, with connectance = 0.12, indicating that 12% of possible viral species-location associations were observed. We found that nestedness = 27.2, reflecting the extent to which viral species detected at less-connected locations represented subsets of those detected at more-connected locations. Finally, network specialisation = 0, indicating no detectable association between particular viral species and sampling locations.

**Figure 5.**
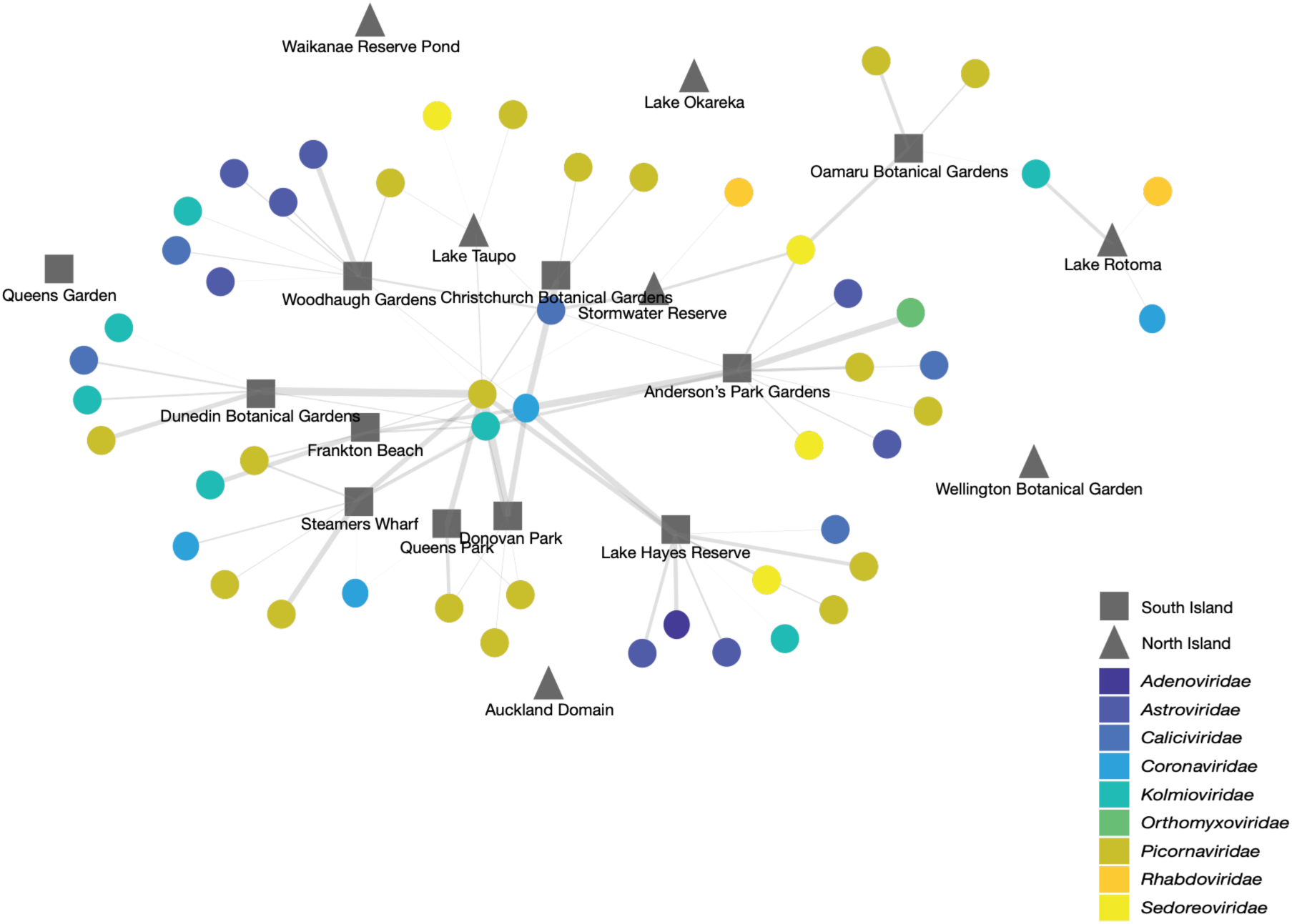
Bipartite network illustrating connectivity between viral species and sampling locations across Aotearoa. Viral species, identified via phylogenetic analysis, are represented by coloured circles, with colours indicating viral family, and sampling locations by grey nodes. Connections indicate detection of a viral species at a sampling location, with connection width proportional to the summed reads per million (RPM) of that viral species for the corresponding sampling location. Five sampling locations were not connected to viral species here since viral sequences detected at these locations could not be resolved beyond the viral family level.

## Discussion

Urban waterfowl occupy an increasingly important interface between wildlife, domestic animals and humans, yet relatively little is known about the diversity and spatial structure of the viruses they harbour (7,22). Using a national scale metatranscriptomic survey, we characterised the faecal viromes of urban waterfowl across Aotearoa, identifying 96 avian viruses spanning 50 species and nine families, including 33 putatively novel virus species. Notably, despite sampling locations spanning over 1,500 km, we found no evidence that geographic distance structured virome composition. Instead, viral species communities were remarkably similar across the country, suggesting that urban waterfowl across Aotearoa harbour broadly similar viral communities.

Ecological communities typically exhibit distance decay, whereby community similarity decreases with increasing geographic separation (49,50). Here, however, we found no association between geography and shared virome community structure. Likewise, neither viral diversity nor overall virome composition differed between the North and South Islands, and virus species were broadly shared among sampling locations. These findings indicate that virome composition is relatively similar among urban waterfowl populations despite substantial geographic separation. Such connectivity likely reflects a combination of local host movements, dispersal among wetlands, shared stopover sites and the widespread distribution of introduced mallards and other waterfowl throughout the country (20). Urban parks may therefore facilitate the movement and persistence of viruses in waterfowl.

Our findings complement previous studies demonstrating that host ecology strongly influences virome composition in wild birds (51–53). While differences in host species, foraging ecology and social behaviour often structure avian viromes (21,22), urban waterfowl appear to represent a particularly well-connected system in which viral exchange occurs readily among locations. The congregation of multiple waterfowl species at artificial ponds, as well as the frequent movement of birds between urban environments, may reduce opportunities for geographic differentiation of viral communities.

This study also substantially expands the known viruses harboured by waterfowl in the Southern Hemisphere. Approximately one-third of the viruses identified represented previously undescribed species, demonstrating that considerable viral diversity remains unsampled even within abundant bird populations (48,53). The greatest diversity was observed within the *Picornaviridae*, where several highly divergent viruses may represent a previously unrecognised genus. Similar patterns of extensive undiscovered diversity have emerged from recent metagenomic surveys of wild birds (48,51–53). In addition, the recovery of a complete influenza virus genome demonstrates the value of sampling urban waterfowl faeces as sentinel hosts for genomic surveillance of avian viruses (54).

Many of the viruses recovered here were closely related to viruses previously described from Australian waterfowl, including those in the gammacoronaviruses, adenoviruses and rotaviruses. Although Aotearoa’s geographic isolation limits natural immigration of waterfowl compared with continental regions, these findings suggest that some viral lineages are shared across the Australasian region. This connectivity may reflect the occasional movement of waterfowl or migratory birds between Australia and Aotearoa, or historical introductions of waterfowl carrying viruses that then persisted and evolved within resident populations. This regional connectivity highlights the importance of considering avian virus ecology at an international scale rather than within national borders alone.

Several limitations of this study should be acknowledged. Primarily, sampling carried out in the lower regions of the South Island was considerably more extensive than that in the Canterbury region and the North Island. Future research could consider a more comprehensive and geographically distributed sampling effort across Aotearoa, which may provide a much better understanding and greater resolution of not only regional but also island patterns in virome composition and connectivity. Secondly, pooling RNA precluded assignment of viruses to individual birds and thus fine-scale analysis of the virus variation among host species. In addition, our sampling focused on urban environments during a single season and future longitudinal studies incorporating repeated sampling across time and non-urban sites would provide a more complete understanding of temporal drivers of virome dynamics and differences in virome composition.

In conclusion, this study provides the first national-scale characterisation of viruses circulating in urban waterfowl across Aotearoa. We demonstrate that these birds harbour diverse viromes, including numerous previously undescribed viruses, while maintaining highly connected viral communities across large geographic distances.

## Data availability

Raw metatranscriptomic sequencing reads generated in this study have been deposited in the NCBI Sequence Read Archive (SRA) under BioProject accession [pending]. Consensus viral genome and polymerase sequences have been deposited in GenBank under accession numbers PZ167563 - PZ167570 and [pending]. All other data supporting the findings of this study are available within the article and its Supplementary Information files.

## Author Contributions

JLG conceptualised the study and acquired the funds. LRH conceptualised the field work and curated the data. LRH, RKF, RMG, JRW, JD, JTT and NSH conducted field work. LRH, JAD and NSH conducted the lab work. LRH, RKF, RMG, SJW and JTT analysed the data and visualised the results. LRH wrote the initial draft. LRH, JLG and SJW wrote the manuscript. All authors read and approved the final manuscript.

## Acknowledgements

We thank S. Heremia, D. Heremia, H. Heremia, T. Heremia and D. Saunders for their invaluable assistance with field sampling and collection of urban waterfowl faecal samples from across Aotearoa. Their support was essential to the successful completion of this study.

## Supplementary Information

**Supplementary Table 1**. Sampling metadata and sequencing information for urban waterfowl faecal libraries. Summary of all pooled metatranscriptomic libraries generated from urban waterfowl faecal samples collected across Aotearoa. Information includes sampling location, collection date, number of individual faecal samples contributing to each pool, sequencing library identifier, total paired-end reads generated, and assembled contigs.

**Supplementary Table 2**. Summary of vertebrate-associated viruses identified in urban waterfowl. Characteristics of the 96 vertebrate-associated viruses identified by metatranscriptomic sequencing. Information includes provisional virus name, viral family and genus, genome type, sampling location(s), closest GenBank match, amino acid sequence identity across the conserved polymerase region (RNA-dependent RNA polymerase or DNA polymerase), GenBank accession number of the closest relative, and whether the virus is considered a putative novel species (<90% amino acid identity to the closest recognised virus).

## References

1. Kessler S, Harder TC, Schwemmle M, Ciminski K. 2021. Influenza A viruses and zoonotic events—are we creating our own reservoirs? Viruses 13(11): 2250. doi:10.3390/v13112250.

2. Chan JFW, To KKW, Chen H, Yuen KY. 2015. Cross-species transmission and emergence of novel viruses from birds. Current Opinion in Virology 10: 63–69. doi:10.1016/j.coviro.2015.01.006.

3. Hess JC, Paré JA. 2004. Viruses of waterfowl. Seminars in Avian and Exotic Pet Medicine 13(4): 176–183. doi:10.1053/j.saep.2004.04.002.

4. Diskin ER, Friedman K, Krauss S, Nolting JM, Poulson RL, Slemons RD, et al. 2020. Subtype diversity of influenza A virus in North American waterfowl: a multidecade study. Journal of Virology 94(11): e02022–19. doi:10.1128/JVI.02022-19.

5. Lebarbenchon C, Stallknecht DE. 2011. Host shifts and molecular evolution of H7 avian influenza virus hemagglutinin. Virology Journal 8: 328. doi:10.1186/1743-422X-8-328.

6. Venkatesh D, Poen MJ, Bestebroer TM, Scheuer RD, Vuong O, Chkhaidze M, et al. 2018. Avian influenza viruses in wild birds: virus evolution in a multihost ecosystem. Journal of Virology 92(15): e00433–18. doi:10.1128/JVI.00433-18.

7. Wille M, Lindqvist K, Muradrasoli S, Olsen B, Järhult JD. 2017. Urbanization and the dynamics of RNA viruses in mallards (Anas platyrhynchos). *Infection*, Genetics and Evolution 51: 89–97. doi:10.1016/j.meegid.2017.03.019.

8. Elmberg J, Berg C, Lerner H, Waldenström J, Hessel R. 2017. Potential disease transmission from wild geese and swans to livestock, poultry and humans: a review of the scientific literature from a One Health perspective. Infection Ecology & Epidemiology 7(1): 1300450. doi:10.1080/20008686.2017.1300450.

9. Owen JC, Hawley DM, Huyvaert KP. 2021. Infectious Disease Ecology of Wild Birds. Oxford University Press, Oxford.

10. National Research Council. 2009. Drivers of zoonotic diseases. In: Keusch GT, Pappaioanou M, Gonzalez MC, Scott KA, Tsai P, editors. Sustaining Global Surveillance and Response to Emerging Zoonotic Diseases. National Academies Press, Washington, DC.

11. Morse SS, Mazet JA, Woolhouse M, Parrish CR, Carroll D, Karesh WB, et al. 2012. Prediction and prevention of the next pandemic zoonosis. The Lancet 380(9857): 1956–1965. doi:10.1016/S0140-6736(12)61684-5.

12. Dennehy JJ. 2017. Evolutionary ecology of virus emergence. Annals of the New York Academy of Sciences 1389(1): 124–146. doi:10.1111/nyas.13304.

13. Jones BA, Grace D, Kock R, Alonso S, Rushton J, Said MY, et al. 2013. Zoonosis emergence linked to agricultural intensification and environmental change. Proceedings of the National Academy of Sciences of the United States of America 110(21): 8399–8404. doi:10.1073/pnas.1208059110.

14. Efsa, ECDC, Alexakis L, Buczkowski H, Ducatez M, Fusaro A, et al. 2025. Avian influenza overview. EFSA Journal 23(1): e9204. doi:10.2903/j.efsa.2025.9204.

15. Enkirch T, Gervelmeyer A, Hallmaier-Wacker L, Melidou A, Willgert K. 2025. Coordinated One Health investigation and management of outbreaks in humans and animals caused by zoonotic avian influenza viruses. EFSA Journal 23(1): e9183. doi:10.2903/j.efsa.2025.9183.

16. Bevins SN, Shriner SA, Cumbee JC, Dilione KE, Douglass KE, Ellis JW, et al. 2022. Intercontinental movement of highly pathogenic avian influenza A(H5N1) clade 2.3.4.4 virus to the United States, 2021. Emerging Infectious Diseases 28(5): 1006–1011. doi:10.3201/eid2805.220318.

17. Blagodatski A, Trutneva K, Glazova O, Mityaeva O, Shevkova L, Kegeles E, et al. 2021. Avian influenza in wild birds and poultry: dissemination pathways, monitoring methods, and virus ecology. Pathogens 10(5): 630. doi:10.3390/pathogens10050630.

18. Greco F, Ravenswater HM, Ruiz-Raya F, D’Avino C, Newell MA, Hewitt J, et al. 2025. Asymptomatic infection and antibody prevalence to co-occurring avian influenza viruses vary substantially between sympatric seabird species following H5N1 outbreaks. Scientific Reports 15: 1435. doi:10.1038/s41598-025-85152-6.

19. Ramey AM, Hill NJ, DeLiberto TJ, Gibbs SEJ, Hopkins MC, Lang AS, et al. 2022. Highly pathogenic avian influenza is an emerging disease threat to wild birds in North America. Journal of Wildlife Management 86(2): e22171. doi:10.1002/jwmg.22171.

20. Stanislawek WL, Tana T, Rawdon TG, Cork SC, Chen K, Fatoyinbo H, et al. 2024. Avian influenza viruses in New Zealand wild birds, with an emphasis on subtypes H5 and H7: their distinctive epidemiology and genomic properties. PLoS One 19(6): e0303756. doi:10.1371/journal.pone.0303756.

21. Wille M, Eden JS, Shi M, Klaassen M, Hurt AC, Holmes EC. 2018. Virus–virus interactions and host ecology are associated with RNA virome structure in wild birds. Molecular Ecology 27(24): 5263–5278. doi:10.1111/mec.14918.

22. Wille M, Shi M, Klaassen M, Hurt AC, Holmes EC. 2019. Virome heterogeneity and connectivity in waterfowl and shorebird communities. ISME Journal 13(10): 2603– 2616. doi:10.1038/s41396-019-0458-0.

23. Gill BJ, West RC. 2018. High densities of water-birds at two New Zealand fresh-water urban lakes. Notornis 65(2): 84. doi:10.63172/353926rdeuzs.

24. French RK, Anderson SH, Cain KE, Greene TC, Minor M, Miskelly CM, et al. 2023. Host phylogeny shapes viral transmission networks in an island ecosystem. Nature Ecology & Evolution 7(11): 1834–1843. doi:10.1038/s41559-023-02192-9.

25. Alley M, Gartrell B. 2019. Wildlife diseases in New Zealand: recent findings and future challenges. New Zealand Veterinary Journal 67(1): 1–11. doi:10.1080/00480169.2018.1520656.

26. Williams M, Gummer H, Powlesland R, Robertson H, Taylor G. 2006. Migrations and movements of birds to New Zealand and surrounding seas. Science for Conservation 232. Department of Conservation, Wellington, New Zealand.

27. Dyer J, Williams M. 2010. An introduction most determined: Mallard (Anas platyrhynchos) to New Zealand. Notornis 57(4): 178–195. doi:10.63172/399767fehrlu.

28. Bolger AM, Lohse M, Usadel B. 2014. Trimmomatic: a flexible trimmer for Illumina sequence data. Bioinformatics 30(15): 2114–2120. doi:10.1093/bioinformatics/btu170.

29. Andrews S. 2010. FastQC: A Quality Control Tool for High Throughput Sequence Data. Babraham Bioinformatics.

30. Li D, Liu C-M, Luo R, Sadakane K, Lam T-W. 2015. MEGAHIT: an ultra-fast single-node solution for large and complex metagenomics assembly via succinct de Bruijn graph. Bioinformatics 31:1674–1676. doi:10.1093/bioinformatics/btv033.

31. Buchfink B, Xie C, Huson DH. 2015. Fast and sensitive protein alignment using DIAMOND. Nature Methods 12(1): 59–60. doi:10.1038/nmeth.3176.

32. Geneious. 2026. Geneious Prime. Biomatters Ltd., Auckland, New Zealand.

33. Langmead B, Salzberg SL. 2012. Fast gapped-read alignment with Bowtie 2. Nature Methods 9:357–359. doi:10.1038/nmeth.1923.

34. Waller SJ, Tortosa P, Thurley T, O’Donnell CFJ, Jackson R, Dennis G, et al. 2024. Virome analysis of New Zealand’s bats reveals cross-species viral transmission among the Coronaviridae. Virus Evolution 10(1): veae008. doi:10.1093/ve/veae008.

35. Nakamura T, Yamada KD, Tomii K, Katoh K. 2018. Parallelization of MAFFT for large-scale multiple sequence alignments. Bioinformatics 34(14): 2490–2492. doi:10.1093/bioinformatics/bty121.

36. Capella-Gutiérrez S, Silla-Martínez JM, Gabaldón T. 2009. trimAl: a tool for automated alignment trimming in large-scale phylogenetic analyses. Bioinformatics 25(15): 1972–1973. doi:10.1093/bioinformatics/btp348.

37. Nguyen LT, Schmidt HA, von Haeseler A, Minh BQ. 2015. IQ-TREE: a fast and effective stochastic algorithm for estimating maximum-likelihood phylogenies. Molecular Biology and Evolution 32(1): 268–274. doi:10.1093/molbev/msu300.

38. Kalyaanamoorthy S, Minh BQ, Wong TKF, von Haeseler A, Jermiin LS. 2017. ModelFinder: fast model selection for accurate phylogenetic estimates. Nature Methods 14(6): 587–589. doi:10.1038/nmeth.4285.

39. Rambaut A. 2018. FigTree version 1.4.4.

40. Oksanen J, Simpson GL, Blanchet FG, Kindt R, Legendre P, Minchin PR, et al. 2025. vegan: Community Ecology Package. R package version 2.6–10. doi:10.32614/CRAN.package.vegan.

41. Google. 2025. Google Earth.

42. Jordano P. ggbipart: an R package for plotting bipartite networks. GitHub repository, pedroj/bipartite_plots.

43. Waller SJ, Butcher RG, Lim L, McInnes K, Holmes EC, Geoghegan JL. 2024. The radiation of New Zealand’s skinks and geckos is associated with distinct viromes. BMC Ecology and Evolution 24: 81. doi:10.1186/s12862-024-02269-4.

44. Dormann CF, Gruber B, Fründ J. 2008. Introducing the bipartite package: analysing ecological networks. R News 8(2): 8–11.

45. Iwamoto M, Shibata Y, Kawasaki J, Kojima S, Li YT, Iwami S, et al. 2021. Identification of novel avian and mammalian deltaviruses provides new insights into deltavirus evolution. Virus Evolution 7(1): veab003. doi:10.1093/ve/veab003.

46. Pinheiro MS, Dias JBL, Petrucci MP, Travassos CEPF, Mendes GS, Santos N. 2023. Molecular characterization of avian rotaviruses F and G detected in Brazilian poultry flocks. Viruses 15(5): 1089. doi:10.3390/v15051089.

47. Vibin J, Chamings A, Klaassen M, Alexandersen S. 2020. Metagenomic characterisation of additional and novel avian viruses from Australian wild ducks. Scientific Reports 10: 22284. doi:10.1038/s41598-020-79413-9.

48. French RK, Filion A, Niebuhr CN, Holmes EC. 2022. Metatranscriptomic comparison of viromes in endemic and introduced passerines in New Zealand. Viruses 14(7): 1364. doi:10.3390/v14071364.

49. Soininen J, McDonald R, Hillebrand H. 2007. The distance decay of similarity in ecological communities. Ecography 30(1): 3–12. doi:10.1111/j.0906-7590.2007.04817.x.

50. Graco-Roza C, Aarnio S, Abrego N, Acosta ATR, Alahuhta J, Altman J, et al. 2022. Distance decay 2.0—a global synthesis of taxonomic and functional turnover in ecological communities. Global Ecology and Biogeography 31(7): 1399–1421. doi:10.1111/geb.13513.

51. Waller SJ, Wierenga JR, Heremia L, Darnley JA, de Vries I, Dubrulle J, et al. 2026. Viral community structure in New Zealand’s aquatic birds is associated with scavenging behaviour. Current Biology, in press.

52. Shan T, Yang S, Wang H, Wang H, Zhang J, Gong G, et al. 2022. Virome in the cloaca of wild and breeding birds revealed a diversity of significant viruses. Microbiome 10: 60. doi:10.1186/s40168-022-01246-7.

53. Lu X, Ji L, Wang H, Zhang Q, Wang X, Liu Y, et al. 2024. Highly diverse RNA viruses and phage sequences concealed within birds. Microbiology Spectrum 12(7): e00802–24. doi:10.1128/spectrum.00802-24.

54. Fornasiero D, Panzarin V, Crimaudo M, Zambon I, Marciano S, Bortolami A, et al. 2026. Integrating wild bird and environmental samples can strengthen avian influenza surveillance. Journal of Applied Ecology. doi:10.1111/1365-2664.70584.

