## Supplementary Table 1 for "Urban waterfowl harbour geographically connected viral communities across Aotearoa New Zealand"

**Supplementary Table 1**. RNA sequencing library RNA concentrations, sampling locations and species sampled.

| **Pooled RNA sequencing library ID** | **City** | **Location** | **Collection Date** | **Species in pool** | **Number of samples in pool** | **Pooled RNA concentration (ng/µL)** | **Total paired-end reads generated** | **No. contigs assembled** |
| --- | --- | --- | --- | --- | --- | --- | --- | --- |
| **APG1** | Invercargill | Andersons Park Gardens | 04.12.2023 | Mallard/Muscovy | 10 | 22.7 | 51,057,935 | 99367 |
| **APG2** | Invercargill | Andersons Park Gardens | 05.12.2023 | Mallard/Muscovy | 10 | 80.5 | 69,120,742 | 266146 |
| **APG3** | Invercargill | Andersons Park Gardens | 26.12.2023 | Mallard/Muscovy | 11 | 108.0 | 87,641,048 | 294008 |
| **AD1** | Auckland | Auckland Domain | 24.11.2023 | Mallard | 7 | 43.4 | 64,093,125 | 193255 |
| **CBG1** | Christchurch | Christchurch Botanical Gardens | 14.11.2023 | Mallard/Paradise | 10 | 38.3 | 81,871,455 | 292417 |
| **WBG1** | Wellington | Wellington Botanical Gardens | 18.11.2023 | Mallard/Paradise | 4 | 37.5 | 101,485,843 | 232407 |
| **DBG1** | Dunedin | Dunedin Botanical Gardens | 02.11.2023 | Mallards | 10 | 46.8 | 126,417,451 | 395534 |
| **DBG2** | Dunedin | Dunedin Botanical Gardens | 20.10.2023 | Mallard | 10 | 21.3 | 67,133,369 | 70314 |
| **DBG3** | Dunedin | Dunedin Botanical Gardens | 11.11.2023 | Mallard | 13 | 34.1 | 72,557,969 | 283075 |
| **DP1** | Invercargill | Donovan Park | 04.12.2023 | Mallard/Muscovy/Pekin | 10 | 11.0 | 73,458,279 | 57341 |
| **DP2** | Invercargill | Donovan Park | 05.12.2023 | Mallard/Muscovy/Pekin | 10 | 63.9 | 80,163,065 | 77082 |
| **DP3** | Invercargill | Donovan Park | 26.12.2023 | Mallard/Muscovy/Pekin/Goose | 10 | 43.5 | 122,737,534 | 342972 |
| **FB1** | Queenstown | Frankton Beach | 18.12.2023 | Mallard/Australian Coot/NZ Scaup | 10 | 83.3 | 104,202,845 | 331868 |
| **FB2** | Queenstown | Frankton Beach | 19.12.2023 | Mallard | 10 | 266.3 | 101,261,544 | 229686 |
| **FB3** | Queenstown | Frankton Beach | 20.12.2023 | Mallard | 10 | 132.4 | 97,709,898 | 248493 |
| **LHR1** | Arrowtown | Lake Hayes Reserve | 18.12.2023 | Mallard/Pekin | 10 | 23.9 | 67,939,636 | 224464 |
| **LHR2** | Arrowtown | Lake Hayes Reserve | 19.12.2023 | Mallard/Pekin/Greylag goose | 10 | 146.3 | 271,773,653 | 498586 |
| **LHR3** | Arrowtown | Lake Hayes Reserve | 20.12.2023 | Mallard/Pekin/Greylag goose | 10 | 113.5 | 71,848,768 | 207037 |
| **LO1** | Rotorua | Lake Okareka | 01.12.2023 | Mallard/Paradise/Goose | 8 | 12.6 | 80,371,271 | 341804 |
| **LR1** | New Plymouth | Lake Rotomanu | 21.11.2023 | Mallard/Paradise | 4 | 6.5 | 74,150,837 | 92318 |
| **LT1** | Taupo | Lake Taupo | 04.12.2023 | Mallard/Paradise | 10 | 25.3 | 93,179,566 | 257931 |
| **LT2** | Taupo | Lake Taupo | 04.12.2023 | Mallard/Paradise | 9 | 63.2 | 83,056,045 | 280766 |
| **OBG1** | Oamaru | Oamaru Botanical Gardens | 07.12.2023 | Mallard/Paradise/Grey Teal | 10 | 29.9 | 77,938,235 | 138442 |
| **OBG2** | Oamaru | Oamaru Botanical Gardens | 08.12.2023 | Mallard/Paradise/Grey Teal/Canadian goose | 10 | 50.4 | 73,210,377 | 229186 |
| **OBG3** | Oamaru | Oamaru Botanical Gardens | 09.12.2023 | Mallard/Grey Teal | 10 | 98.0 | 70,056,699 | 304453 |
| **QG1** | Nelson | Queens Garden | 28 and 30.12.2023 | Mallard/Goose | 8 | 32.7 | 82,833,401 | 73635 |
| **QP1** | Invercargill | Queens Park | 04.12.2023 | Mallard | 10 | 17.8 | 35,908,646 | 48963 |
| **QP2** | Invercargill | Queens Park | 05.12.2023 | Mallard | 10 | 39.6 | 74,042,537 | 148649 |
| **QP3** | Invercargill | Queens Park | 26.12.2023 | Mallard | 10 | 62.8 | 104,085,243 | 286989 |
| **SW1** | Queenstown | Steamers Wharf | 19.12.2023 | Mallard | 10 | 21.3 | 99,578,754 | 201449 |
| **SW2** | Queenstown | Steamers Wharf | 20.12.2023 | Mallard | 10 | 68.6 | 51,947,442 | 70726 |
| **SW3** | Queenstown | Steamers Wharf | 21.12.2023 | Mallard/Paradise | 10 | 52.3 | 58,206,048 | 224956 |
| **SR1** | Papamoa | Stormwater reserve | 30.11.2023 | Mallard/Paradise | 10 | 20 | 80,861,626 | 312966 |
| **WRP1** | Paraparaumu | Waikanae reserve pond | 19.11.2023 | Black Swan | 7 | 12 | 78,826,492 | 72677 |
| **WH1** | Dunedin | Woodhaugh Gardens | 24.10.23 and 11.12.2023 | Mallard/Pekin | 11 | 98.3 | 87,145,375 | 186314 |
| **WH2** | Dunedin | Woodhaugh Gardens | 12.12.2023 | Mallard/Pekin | 10 | 101.3 | 65,785,565 | 269406 |
| **WH3** | Dunedin | Woodhaugh Gardens | 13.12.2023 | Mallard/Pekin | 10 | 154.3 | 109,846,394 | 350703 |
