## Supplementary Table 2 for "Urban waterfowl harbour geographically connected viral communities across Aotearoa New Zealand"

**Supplementary Table 2**. Summary of vertebrate-associated viruses identified in urban waterfowl. Characteristics of the 96 vertebrate-associated viruses identified by metatranscriptomic sequencing. Information includes provisional virus name, viral family, sampling location(s), amino acid sequence identity across the conserved polymerase region (RNA-dependent RNA polymerase or DNA polymerase), GenBank accession number of the closest relative, and whether the virus is considered a putative novel species (<90% amino acid identity to the closest recognised virus).

| **Contig ID** | **Library ID** | **Virus Family** | **Location Sampled** | **Virus Name** | **Sequence Length (aa)** | **Closest genetic relative in GenBank** | **Percentage amino acid identity (%)** | **Novel virus species** |
| --- | --- | --- | --- | --- | --- | --- | --- | --- |
| k141_242556 | LHR3 | *Adenoviridae* | Lake Hayes Reserve | Duck aviadenovirus | 442 | QQD36933.1 | 99.77% | No |
| k141_194195 | APG3 | *Astroviridae* | Anderson Park Gardens | Waterfowl astrovirus APG3.2 | 352 | AMO03288.1 | 88.51% | Yes |
| k141_299416 | APG3 | *Astroviridae* | Anderson Park Gardens | Waterfowl astrovirus APG3.1 | 160 | QCR98314.1 | 82.71% | Yes |
| k141_248920 | LHR3 | *Astroviridae* | Lake Hayes Reserve | Duck astrovirus | 437 | AID55206.1 | 92.23% | No |
| k141_449943 | LHR2 | *Astroviridae* | Lake Hayes Reserve | Duck astrovirus | 223 | WPX56601.1 | 92.38% | No |
| k141_239345 | LHR3 | *Astroviridae* | Lake Hayes Reserve | Duck astrovirus | 106 | ADB79803.1 | 96.23% | No |
| k141_50557 | LHR3 | *Astroviridae* | Lake Hayes Reserve | Duck astrovirus | 197 | AUX13653.1 | 97.97% | No |
| k141_213705 | WH2 | *Astroviridae* | Woodhaugh Gardens | Waterfowl astrovirus WH2.1 | 160 | AHX26591.1 | 88.75% | Yes |
| k141_129769 | WH2 | *Astroviridae* | Woodhaugh Gardens | Duck astrovirus | 151 | AHX26591.1 | 96.03% | No |
| k141_56261 | WH1 | *Astroviridae* | Woodhaugh Gardens | Duck astrovirus | 247 | QQD37002.1 | 92.44% | No |
| k141_363694 | WH3 | *Astroviridae* | Woodhaugh Gardens | Waterfowl astrovirus WH3.1 | 449 | QQD37003.1 | 89.63% | Yes |
| k141_277610 | WH2 | *Caliciviridae* | Woodhaugh Gardens | Duck calicivirus | 340 | QKN88796.1 | 96.76% | No |
| k141_38642 | SR1 | *Caliciviridae* | Stormwater Reserve | Duck calicivirus | 1407 | WYV83164.1 | 96.96% | No |
| k141_31174 | APG1 | *Caliciviridae* | Anderson Park Gardens | Duck calicivirus | 129 | QEG79135.1 | 94.57% | No |
| k141_125742 | LT2 | *Caliciviridae* | Lake Taupō | Duck calicivirus | 246 | WYV83163.1 | 95.53% | No |
| k141_341994 | WH3 | *Caliciviridae* | Woodhaugh Gardens | Duck calicivirus | 575 | QEG79135.1 | 95.48% | No |
| k141_82639 | DBG3 | *Caliciviridae* | Dunedin Botanical Gardens | Duck calicivirus | 445 | WYV95937.1 | 99.78% | No |
| k141_37124 | DBG1 | *Caliciviridae* | Dunedin Botanical Gardens | Duck calicivirus | 296 | WYV95937.1 | 99.32% | No |
| k141_124812 | APG3 | *Caliciviridae* | Anderson Park Gardens | Waterfowl calicivirus APG3.1 | 271 | AXF38742.1 | 47.72% | Yes |
| k141_95066 | LHR3 | *Caliciviridae* | Lake Hayes Reserve | Waterfowl calicivirus LHR3.1 | 144 | AXF38742.1 | 51.33% | Yes |
| k141_199837 | LHR3 | *Coronaviridae* | Lake Hayes Reserve | Duck gammacoronavirus | 482 | QDY92334.1 | 99.17% | No |
| k141_155704 | SW3 | *Coronaviridae* | Steamers Wharf | Duck gammacoronavirus | 501 | QDY92334.1 | 96.81% | No |
| k141_179826 | QP3 | *Coronaviridae* | Queens Park | Duck gammacoronavirus | 106 | WMM65745.1 | 100.00% | No |
| k141_149800 | WH2 | *Coronaviridae* | Woodhaugh Gardens | Duck gammacoronavirus | 146 | QDY92358.1 | 100.00% | No |
| k141_77467 | LR1 | *Coronaviridae* | Lake Rotomanu | Duck gammacoronavirus | 142 | AKQ98494.1 | 98.59% | No |
| k141_117142 | APG2 | *Coronaviridae* | Anderson Park Gardens | Duck gammacoronavirus | 2691 | QDY92334.1 | 99.00% | No |
| k141_183883 | SW1 | *Coronaviridae* | Steamers Wharf | Duck gammacoronavirus | 110 | AWR88348.1 | 100.00% | No |
| k141_221937 | LHR1 | *Coronaviridae* | Lake Hayes Reserve | Duck gammacoronavirus | 781 | QDY92358.1 | 99.74% | No |
| k141_532479 | LHR2 | *Coronaviridae* | Lake Hayes Reserve | Duck gammacoronavirus | 2697 | QDY92358.1 | 99.22% | No |
| k141_340684 | DP3 | *Coronaviridae* | Donovan Park | Duck gammacoronavirus | 2691 | QDY92358.1 | 99.18% | No |
| k141_5295 | CBG1 | *Coronaviridae* | Christchurch Botanical Gardens | Duck gammacoronavirus | 2691 | QDY92358.1 | 99.22% | No |
| k141_77171 | SW2 | *Coronaviridae* | Steamers Wharf | Duck gammacoronavirus | 124 | XDG24379.1 | 96.77% | No |
| k141_278682 | APG3 | *Coronaviridae* | Anderson Park Gardens | Duck gammacoronavirus | 134 | QEG79146.1 | 96.27% | No |
| k141_310568 | FB1 | *Coronaviridae* | Frankton Beach | Duck gammacoronavirus | 1115 | XDG24341.1 | 98.46% | No |
| k141_66756 | DP2 | *Coronaviridae* | Donovan Park | Duck gammacoronavirus | 252 | XDG24370.1 | 97.62% | No |
| k141_252195 | FB2 | *Kolmioviridae* | Frankton Beach | Duck dalvirus | 220 | YP_009553679.1 | 98.38% | No |
| k141_267998 | FB3 | *Kolmioviridae* | Frankton Beach | Duck dalvirus | 247 | YP_009553679.1 | 98.38% | No |
| k141_86774 | APG1 | *Kolmioviridae* | Anderson Park Gardens | Duck dalvirus | 232 | YP_009553679.1 | 98.92% | No |
| k141_73670 | DP2 | *Kolmioviridae* | Donovan Park | Duck dalvirus | 220 | YP_009553679.1 | 99.46% | No |
| k141_490780 | LHR2 | *Kolmioviridae* | Lake Hayes Reserve | Waterfowl kolmiovirus LHR2.1 | 144 | YP_009553679.1 | 88.89% | Yes |
| k141_440322 | DBG1 | *Kolmioviridae* | Dunedin Botanical Gardens | Waterfowl kolmiovirus DBG1.1 | 105 | YP_009553679.1 | 87.62% | Yes |
| k141_10619 | WH1 | *Kolmioviridae* | Woodhaugh Gardens | Waterfowl kolmiovirus WH1.1 | 123 | YP_009553679.1 | 89.43% | Yes |
| k141_85071 | SW3 | *Kolmioviridae* | Steamers Wharf | Duck dalvirus | 141 | YP_009553679.1 | 90.07% | No |
| k141_16386 | DBG2 | *Kolmioviridae* | Dunedin Botanical Gardens | Waterfowl kolmiovirus DBG2.1 | 128 | YP_009553679.1 | 89.06% | Yes |
| k141_74540 | DP3 | *Kolmioviridae* | Donovan Park | Duck dalvirus | 170 | YP_009553679.1 | 90.59% | No |
| k141_366368 | FB1 | *Kolmioviridae* | Frankton Beach | Waterfowl kolmiovirus FB1.1 | 185 | YP_009553679.1 | 87.57% | Yes |
| k141_179725 | OBG2 | *Kolmioviridae* | Ōamaru Botanical Gardens | Duck dalvirus | 104 | FAA01385.1 | 100.00% | No |
| k141_121277 | LR1 | *Kolmioviridae* | Lake Rotomanu | Duck dalvirus | 226 | FAA01385.1 | 99.49% | No |
| k141_18875 | APG3 | *Orthomyxoviridae* | Anderson Park Gardens | Influenza A virus | 757 | ADF45464.1 | 100.00% | No |
| k141_109625 | LR1 | *Rhabdoviridae* | Lake Rotomanu | Waterfowl rhabdovirus LR1.1 | 129 | AXF38662.1 | 74.22% | Yes |
| k141_18115 | SR1 | *Rhabdoviridae* | Stormwater Reserve | Waterfowl rhabdovirus SR1.1 | 80 | YP_009094346.1 | 51.92% | Yes |
| k141_304124 | SR1 | *Rhabdoviridae* | Stormwater Reserve | Waterfowl rhabdovirus SR1.2 | 123 | AXF38754.1 | 74.80% | Yes |
| k141_8925 | OBG3 | *Sedoreoviridae* | Ōamaru Botanical Gardens | Duck Rotavirus G | 302 | UAJ21467.1 | 98.34% | No |
| k141_46491 | APG1 | *Sedoreoviridae* | Anderson Park Gardens | Duck Rotavirus G | 144 | UAJ21467.1 | 97.92% | No |
| k141_161731 | SR1 | *Sedoreoviridae* | Stormwater Reserve | Duck Rotavirus G | 319 | UAJ21467.1 | 98.75% | No |
| k141_49668 | SW2 | *Sedoreoviridae* | Steamers Wharf | Duck Rotavirus G | 108 | UAJ21467.1 | 100.00% | No |
| k141_98258 | APG2 | *Sedoreoviridae* | Anderson Park Gardens | Duck Rotavirus G | 484 | QQD36988.1 | 96.28% | No |
| k141_77169 | LHR1 | *Sedoreoviridae* | Lake Hayes Reserve | Duck Rotavirus F | 167 | WIF29857.1 | 97.01% | No |
| k141_138728 | LT1 | *Sedoreoviridae* | Lake Taupo | Duck Rotavirus F | 101 | USL85483.1 | 98.86% | No |
| k141_100373 | APG3 | *Picornaviridae* | Anderson Park Gardens | Waterfowl picornavirus APG3.1 | 1749 | QUS52520.1 | 89.72% | Yes |
| k141_107193 | LHR3 | *Picornaviridae* | Lake Hayes Reserve | Megrivirus | 2868 | QMI57985.1 | 93.11% | No |
| k141_11592 | OBG1 | *Picornaviridae* | Ōamaru Botanical Gardens | Waterfowl megrivirus OBG1.1 | 953 | QUS52520.1 | 89.72% | Yes |
| k141_12869 | DP1 | *Picornaviridae* | Donovan Park | Megrivirus | 1523 | QDY92331.1 | 98.13% | No |
| k141_131135 | CBG1 | *Picornaviridae* | Christchurch Botanical Gardens | Waterfowl hepatovirus CBG1.1 | 357 | WKV33068.1 | 55.63% | Yes |
| k141_140328 | WH1 | *Picornaviridae* | Woodhaugh Gardens | Megrivirus | 118 | QMI57984.1 | 98.31% | No |
| k141_150098 | QP3 | *Picornaviridae* | Queens Park | Waterfowl megrivirus QP3.1 | 692 | XBC28235.1 | 75.15% | Yes |
| k141_152586 | DBG3 | *Picornaviridae* | Dunedin Botanical Gardens | Waterfowl passerivirus DBG3.1 | 1548 | QOQ52505.1 | 36.63% | Yes |
| k141_157133 | LHR1 | *Picornaviridae* | Lake Hayes Reserve | Waterfowl picornavirus LHR1.1 | 260 | WGL41079.1 | 53.28% | Yes |
| k141_168848 | SW1 | *Picornaviridae* | Steamers Wharf | Waterfowl sapelovirus SW1.1 | 191 | QUS52757.1 | 66% | Yes |
| k141_18567 | DBG2 | *Picornaviridae* | Dunedin Botanical Gardens | Megrivirus | 2815 | QMI57985.1 | 98.24% | No |
| k141_187970 | FB2 | *Picornaviridae* | Frankton Beach | Megrivirus A | 765 | QUS52520.1 | 93.59% | No |
| k141_200451 | DBG3 | *Picornaviridae* | Dunedin Botanical Gardens | Waterfowl megrivirus DBG3.1 | 2597 | QMI57991.1 | 82.31% | Yes |
| k141_214226 | SW1 | *Picornaviridae* | Steamers Wharf | Megrivirus A | 785 | QUS52520.1 | 93.63% | No |
| k141_21815 | SW2 | *Picornaviridae* | Steamers Wharf | Megrivirus | 2836 | QMI57985.1 | 97.73% | No |
| k141_225716 | CBG1 | *Picornaviridae* | Christchurch Botanical Gardens | Megrivirus | 713 | QMI57985.1 | 98.18% | No |
| k141_226833 | FB3 | *Picornaviridae* | Frankton Beach | Megrivirus | 392 | QDY92331.1 | 98.21% | No |
| k141_23042 | OBG2 | *Picornaviridae* | Ōamaru Botanical Gardens | Waterfowl passerivirus OBG2.1 | 1223 | AVN87703.1 | 36.91% | Yes |
| k141_235905 | LT2 | *Picornaviridae* | Lake Taupo | Megrivirus | 200 | QMI57984.1 | 99.50% | No |
| k141_236741 | OBG2 | *Picornaviridae* | Ōamaru Botanical Gardens | Waterfowl passerivirus OBG2.2 | 319 | WPS63802.1 | 47.76% | Yes |
| k141_279766 | LT2 | *Picornaviridae* | Lake Taupo | Waterfowl aalivirus LT2.1 | 148 | YP_009026377.1 | 61.64% | Yes |
| k141_284173 | OBG3 | *Picornaviridae* | Ōamaru Botanical Gardens | Waterfowl passerivirus OBG3.1 | 303 | WPS63802.1 | 47.74% | Yes |
| k141_321024 | OBG3 | *Picornaviridae* | Ōamaru Botanical Gardens | Waterfowl sicinivirus OBG3.1 | 296 | QJE50373.1 | 68.58% | Yes |
| k141_3220 | QP1 | *Picornaviridae* | Queens Park | Megrivirus | 890 | QMI57984.1 | 99.13% | No |
| k141_323802 | DP3 | *Picornaviridae* | Donovan Park | Ludopivirus A1 | 281 | QUS52516.1 | 96.44% | No |
| k141_327238 | DP3 | *Picornaviridae* | Donovan Park | Megrivirus | 822 | QDY92331.1 | 97.98% | No |
| k141_37835 | APG1 | *Picornaviridae* | Anderson Park Gardens | Waterfowl megrivirus APG1.1 | 761 | QUS52520.1 | 88.70% | Yes |
| k141_383827 | DP3 | *Picornaviridae* | Donovan Park | Anativirus-like virus | 422 | QMI57979.1 | 92.31% | No |
| k141_45843 | DP2 | *Picornaviridae* | Donovan Park | Waterfowl megrivirus DP2.1 | 316 | QUS52521.1 | 79.05% | Yes |
| k141_472761 | LHR2 | *Picornaviridae* | Lake Hayes Reserve | Megrivirus | 146 | QDY92331.1 | 98.63% | No |
| k141_548745 | LHR2 | *Picornaviridae* | Lake Hayes Reserve | Megrivirus | 599 | QMI57984.1 | 98.50% | No |
| k141_56168 | DBG1 | *Picornaviridae* | Dunedin Botanical Gardens | Megrivirus | 598 | QDY92331.1 | 98.16% | No |
| k141_71168 | SW2 | *Picornaviridae* | Steamers Wharf | Waterfowl sapelovirus SW2.1 | 2147 | QDY92367.1 | 84.47% | Yes |
| k141_76592 | SW2 | *Picornaviridae* | Steamers Wharf | Waterfowl sapelovirus SW2.1 | 2508 | QUS52756.1 | 60.64% | Yes |
| k141_85902 | LHR3 | *Picornaviridae* | Lake Hayes Reserve | Waterfowl hepatovirus LHR3.1 | 340 | QUS52647.1 | 43.66% | Yes |
| k141_87065 | CBG1 | *Picornaviridae* | Christchurch Botanical Gardens | Waterfowl anativirus CBG1.1 | 285 | QMI57979.1 | 89.12% | Yes |
| k141_83211 | APG2 | *Picornaviridae* | Anderson Park Gardens | Anativirus A1 APG2.1 | 192 | YP_164335.1 | 73.30% | Yes |
| k141_99644 | QP2 | *Picornaviridae* | Queens Park | Megrivirus | 140 | QMI57984.1 | 99.29% | No |
